# Transcriptome analysis of the zebrafish *atoh7−/−* mutant, *lakritz*, highlights Atoh7-dependent genetic networks with potential implications for human eye diseases

**DOI:** 10.1101/2020.04.09.033704

**Authors:** Giuseppina Covello, Fernando J. Rossello, Michele Filosi, Felipe Gajardo, Anne-Laure Duchemin, Beatrice F. Tremonti, Michael Eichenlaub, Jose M. Polo, David Powell, John Ngai, Miguel L. Allende, Enrico Domenici, Mirana Ramialison, Lucia Poggi

## Abstract

Expression of the bHLH transcription protein Atoh7 is a crucial factor conferring competence to retinal progenitor cells for the development of retinal ganglion cells. A number of studies have emerged establishing *ATOH7* as a retinal disease gene. Remarkably, such studies uncovered *ATOH7* variants associated with global eye defects including optic nerve hypoplasia, microphthalmia, retinal vascular disorders and glaucoma. The complex genetic networks and cellular decisions arising downstream of *atoh7* expression, and how their dysregulation cause development of such disease traits remains unknown. To begin to understand such Atoh7-dependent events *in vivo* we performed transcriptome analysis of wild type and *atoh7* mutant (*lakritz*) zebrafish embryos at the onset of retinal ganglion cell differentiation. We investigated *in silico* interplays of *atoh7* and other disease-related genes and pathways. By network reconstruction analysis of differentially expressed genes we identified gene clusters enriched in retinal development, cell cycle, chromatin remodelling, stress response and Wnt pathways. By weighted gene coexpression network we identified coexpression modules affected by the mutation and enriched in retina development genes tightly connected to *atoh7*. We established the groundwork whereby Atoh7-linked cellular and molecular processes can be investigated in the dynamic multi-tissue environment of the developing normal and diseased vertebrate eye.

## INTRODUCTION

Retinal ganglion cells (RGCs) collect visual information from the neural retina in the eye and convey it to the visual cortex of the brain. In healthy people, this information is transmitted along the optic nerve, which is composed mainly of axons formed from the cell bodies of RGCs. Inherited diseases affecting the development of RGCs and the optic nerve can interrupt this information flow causing permanent blindness ^1,2^. Atoh7 is an evolutionarily conserved, developmentally regulated transcription factor crucial for the genesis of RGCs in different vertebrate models ^3–7^. Studies have shown that induced or naturally occurring mutations in the *atoh7* gene result in retinal progenitor cells (RPCs) failing to develop into RGCs and the optic nerve ^3,5,6^. This occurs concomitantly with a comparable increase in the other main retinal cell types namely, amacrine, horizontal, bipolar, photoreceptor and Müller glial cells, suggesting a fate switch in the RPCs ^3–6,8–11^. An increasing number of studies highlight *ATOH7* as an emerging candidate for eye diseases in humans. Variations in the *ATOH7* locus have been associated with optic nerve hypoplasia (ONH) and aplasia (ONA) ^12–15^, further pointing towards a crucial role of *atoh7* in RGC genesis and optic nerve development. Remarkably, a number of studies also have emerged, which highlight *ATOH7* variants as associated with multiple eye disease traits. For example, genome-wide association studies have reported *ATOH7* variants linked to glaucoma-related traits ^16–23^. Likewise, multiple global eye developmental defects causing congenital blindness have been associated with mutations in *ATOH7*; these include autosomal recessive congenital disorders of the retinal vasculature, such as retinal non-attachment (NCRNA) and persistent hyperplastic primary vitreous (PHPV)(OMIM:# 221900, ORPHA:91495) ^12,14,24–29^ as well as corneal opacity, microcornea and microphthalmia (ORPHA:289499) ^14,30^. Whilst the development of such global eye disorders might result from the association of variations in *ATOH7* and other genes ^20,31,32^, these findings suggest direct or indirect requirements of Atoh7 in multiple molecular and cellular interactions during ocular tissue development. For example, retinal vascular disorders likely result from the interruption of RGC development by the *ATOH7* mutations ^13,28^. The Atoh7-regulated gene networks involved, and how their disruption contribute to the interruption of these retinal neural-vascular interactions remain unknown.

Given the well-established requirement of Atoh7 for RGC development in vertebrate models, several studies from different species have emerged to identify Atoh7-dependent gene batteries; which may also serve as instructive reprogramming factors to efficiently and irreversibly direct retinal progenitor or stem cells towards RGC differentiation pathways ^7,33–42^. Whilst considerable progress has been made in this direction, whether Atoh7 is a master regulator instructing RGC differentiation programs remains debatable. For example, Atoh7 forced expression is often insufficient for bursting RGC fate commitment ^42–44^. Investigators have also shown that, similarly to other bHLH factors in the developing central nervous system ^45^, *atoh7* expression levels are found in cycling retinal progenitor cells acquiring multiple retinal fates ^8–10,46–49^. Concordantly, an increasing number of evidences suggest Atoh7-requirement in the control of retinal progenitor cell cycle progression as well as their competence and RGC identity acquisition ^47,50–54^. Furthermore, *atoh7* expression is transient, being turned on just before the last mitotic division of RGC progenitors and downregulated shortly after the terminal division of the specified RGC daughter ^8,48^. All of these observations call for the need of a deeper understanding as to the cell context- and *atoh7*-dependent regulatory networks that integrate multipotency, self-renewal, lineage-restriction, and cell-specific interactions immediately preceding and during RGC and optic nerve genesis. Also important is to investigate these networks *in vivo*, while observing integrated cell behaviours and vascular-neural interactions occurring in the physiological environment of the developing eye. This approach should inform on how, deregulation of key genes and molecular pathways might affect eye tissue development and interactions, thereby potentially contributing to the described *ATOH7*-associated global eye developmental disorders ^55^.

To this end, the zebrafish has long been valued as a paradigm for disentangling the genetics and cell biology of fundamental eye developmental processes ^56,57^. The rapidly and externally developing transparent zebrafish embryos are amenable to easy genetic manipulation, thus allowing fast generation and identification of mutants modelling human ocular genetic disorders ^58–64^. Such disease models can be concurrently investigated in large-scale genetics, drug screening, *in vivo* cell biology of early disease development as well as behavioural assays ^65–68^. These potentials substantially aid fast progress in the validation of human genome association studies and in preclinical therapy development paths towards the early diagnosis and/or restoration of visual function ^69–72^.

We here begin to explore the potentials of the *lakritz* zebrafish mutant carrying a loss of function mutation in the *atoh7* gene ^6^ to investigate Atoh7-regulated gene networks and interrogate how deregulation of these networks during early onset of RGC genesis might contribute to the development of *atoh7*-associated eye disorders. With the analysis of available microarray data, we provide a cohort of statistically significantly regulated Atoh7 target genes, including previously known Atoh7-targets such as *atoh7* itself ^33,53^. Remarkably, at this early RGC developmental time-point, the most significant targets comprehend previously unreported eye field transcription factors, Wnt signalling pathway components, chromatin and cytoskeletal regulators, and even stress-response proteins as major Atoh7-regulated genes. Interestingly, components of these pathways include eye disease gene markers.

With these data in hand, we can now begin to exploit the power of zebrafish as an *in vivo* vertebrate model to assess how dysregulation of one or more of these components might together affect the developing native ocular tissues. Understanding the cellular context and dynamics of these Atoh7-dependent networks will hopefully provide us with a next step forward in the identification of potential targets for the early detection and/or specific treatment of inherited eye diseases such as retinal-vascular disorders.

## MATERIALS AND METHODS

### Wild-Type and Transgenic Zebrafish

Fish used in this study were identified heterozygous carriers of the *lakritz* mutation ^6^ crossed in the (AB/AB) background as well as transgenic *tg(lakritz/atoh7:gap43-RFP)* heterozygous carriers ^9,53^. All fish were maintained at 26-28°C and embryos raised at 28.5°C or 32°C and staged as described previously ^73,74^. Embryos were obtained by breeding adult male and female fish at ratio 1:3. After fertilization, eggs were collected and maintained at 28.5°C and staged using standard morphological criteria ^73^. Fish were kept and experiments were performed in accordance to local animal welfare agencies and European Union animal welfare guidelines.

### Eyes and Body sample collection

Single pairs of eyes were dissected from single embryos at 25, 28, 35, 48, 72, and 96 hpf stages and snap-frozen in liquid nitrogen, and stored at −80 °C. Embryos older than 28 hpf were first anesthetized for 5–10 min in Ethyl 3-aminobenzoate methanesulfonate (MS-222) (Sigma-Aldrich, Saint Louis, MO, USA) in E3 medium. The corresponding body of each embryo was collected and used immediately to perform genotyping analysis to identify the corresponding to *lakritz* and wild type eyes. All embryos were collected from the same batches of fish stock to maintain a uniform genetic background.

### DNA extraction from zebrafish body biopsies and genotyping

Genomic DNA extraction from each single body was performed in 100 μl of lysis buffer containing Proteinase K-20mg/ml (EuroClone S.p.A. Milan, Italy), 2 M Tris-HCl ph 8.0, 0.5 M EDTA ph 8.0 and 5 M NaCl, 20 % SDS in a final volume of 50 μl ultra H_2_0. After 3 hours of incubation at 65°C, the gDNA was purified with an Ethanol precipitation step and resuspended in 50 μl of Dnase/Rnase H_2_O. The genotyping was performed by Restriction Fragment Length Polymorphism (RFLP) assay as previously described ^6^. An ~ 300 bp fragment of *atoh7* was PCR amplified with 1 U Taq DNA polymerase (Applied Biosystems by Life Technologies, Carlsbad, CA, USA) according to manufacturer protocols in a 30μl PCR mix containing 100ng of purified gDNA (from each single embryo body) with the following primers: Forward 5′ CCGGAATTACATCCCAAGAAC-3′ and Reverse 5′-GGCCATGATGTAGCTCAGAG-3′. The PCR conditions were as follows: initial denaturation (95°C for 5 minutes), followed by 40 cycles of denaturation (95°C for 45 seconds), annealing (56°C for 45 seconds), extension (72°C for45 second), and a final extension at 72 °C for 5 minutes. The resulting PCR product was digested with StuI restriction enzyme (NEB, New England Biolabs, Ipswich, MA, USA), according to manufacturer protocols. The digested product was analysed on a 2% agarose gel in 1X Tris-Acetate EDTA (TAE) buffer (Sigma-Aldrich, Saint Louis, MO, USA) to highlight wt or *lakritz* mutant corresponding fragments. The L44P mutation ^6^eliminates a restriction site found in the published L44 allele The L44P mutation eliminates a restriction site found in the published L44 allele (Masai et al., 2000) and therefore can be visualised as an undigested ~ 300 bp fragment rather than the ~ 100 and ~ 200 bp fragments expected from the wt condition (**Fig. 1**). 1Kb DNA Ladder was used as a reference (Gene ruler 1Kb plus, Thermo Scientific, Carlsbad, CA, USA).

**Figure 1:**
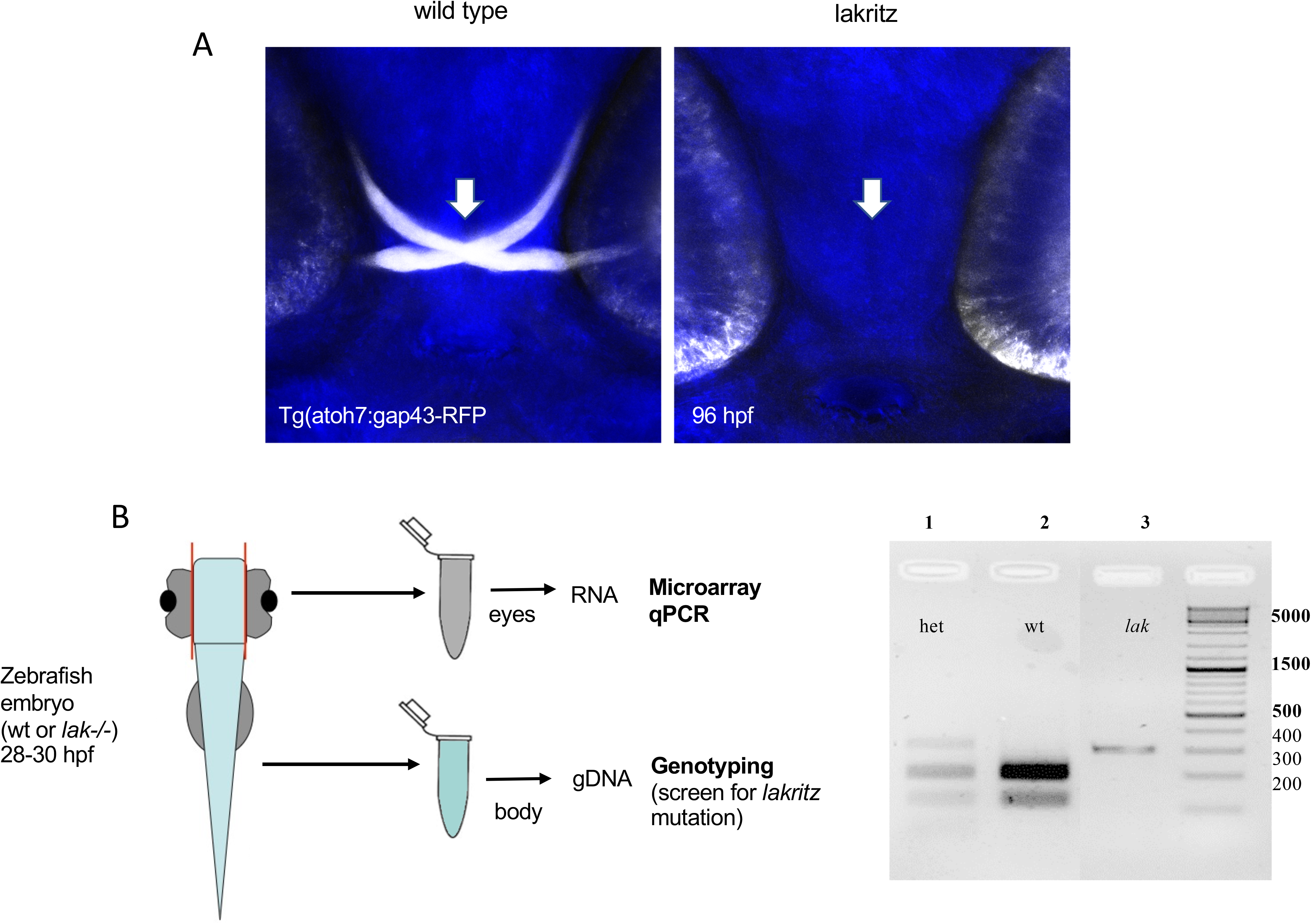
Scheme of the experimental design for the comparative marray analysis. *A)* Confocal images showing examples of wild type and *lak−/−* (*lakritz*); *tg(atoh7:gap43-RFP)* embryos at 96 hpf. The RFP-positive optic chiasm and RGCs are absent in the retina of a *lakritz* embryo. *B)* Pairs of eyes were dissected from single embryos at 28-30 hpf embryos. Genotyping on the gDNA extracted from each corresponding cell body was performed to identify *lakritz* and wt embryos (see materials and methods section). The RNA extracted from each pair of eyes corresponding to either a *lakritz* or wt embryo was amplified and used for the microarray analysis and qPCR expression analysis.

### Affymetrix arrays hybridization and analysis

For the microarray analysis, 3 pairs of wt and *lakritz* 28-30 hpf embryos representing 3 biological replicates were dissected and placed in Trizol reagent (Thermo scientific Life Technologies, Carlsbad, CA, USA) for the total RNA extraction according to manufacturer’s instructions. T7-based linear amplification of the mRNA was performed using the megascript kit from Ambion. Hybridisation was performed on the Affymetrix GeneChip platform and processed according to standard procedure ^75^.

Correspondence between Zebrafish Affymetrix probesets and EnsEMBL gene annotations were retrieved using BioMart (EnsEMBL Version 84, March 2016) ^76^. Batch effect removal was applied to adjust for known batch effect by first filtering the normalized matrix of intensities discarding probes with total abundance between samples lower than first quartile (Q1 = 16.26), and then correcting intensities using ComBat package ^77^ with extraction day as the known batch **(Supp. Fig. 1)**. Differential expression analysis between mutant and wild-type samples was performed using Limma package ^78^For probes showing statistically significant differential expression (adj. P-value < 0.05), annotations of corresponding genes were retrieved from the EnsEMBL database using BiomaRt package ^79^.

### Quantitative Real-Time PCR (qRT-PCR)

For the qRT-PCR analysis of *anillin* and *atoh7*, total RNA from five pulled pairs of frozen eyes, corresponding to either *lakritz* or wild type embryos, was used. After Turbo DNase treatment (Thermo scientific-Ambion, Carlsbad, CA, USA), according to manufacturer instructions, the RNA concentrations were measured with a Nanodrop ND-1000 spectrophotometer (NanoDrop Technologies Inc., Wilmington, USA). The RNA integrity was verified by loading the samples on 1% agarose gel that was run at 100 Volts in TBE 1X. 500 ng of extracted RNA from each sample was restrotrascribed with a RevertAidTM First Strand cDNA Synthesis Kit (Thermo SCIENTIFIC, Carlsbad, CA, USA), following manufacturer’s protocol. Quantitative Real-time PCR reactions were performed on a Bio-Rad CFX96 Termo-cycler with Kapa Syber Fast qPCR master mix (2x) kit (Sigma-Aldrich, Saint Louis, MO, USA), according to the manufacturer’s instructions. Templates were 1:10 diluted cDNA samples. For the negative controls cDNAs were replaced by DEPC water. All real-time assays were carried using 10ng of cDNA. The PCR profile was: 15 seconds at 95°C, followed by 40 cycles 60° C for 20 seconds, 72° C for 40 seconds. For the melting curve, 0.5°C was increased every 5 s from 65°C to 95 °C. All reactions were run in triplicate and both glyceraldehyde-3-phosphate dehydrogenase (GAPDH) and Ubiquitin Conjugating Enzyme E2 A (UBE2A) were used as reference genes. Each experiment was performed in triplicate and repeated two times. The relative gene expression was calculated using the ΔΔCT method. Statistical analyses were performed with Prism 5 (GraphPad Software, San Diego, CA), and statistical significance was set to P<0.05 for all experiments. The values are expressed as mean±SEM, and the differences between groups were investigated using unpaired two-tailed Student t-test (GraphPad Software, San Diego, CA). A list of primers can be found in **Table 1**.

**Table 1.**
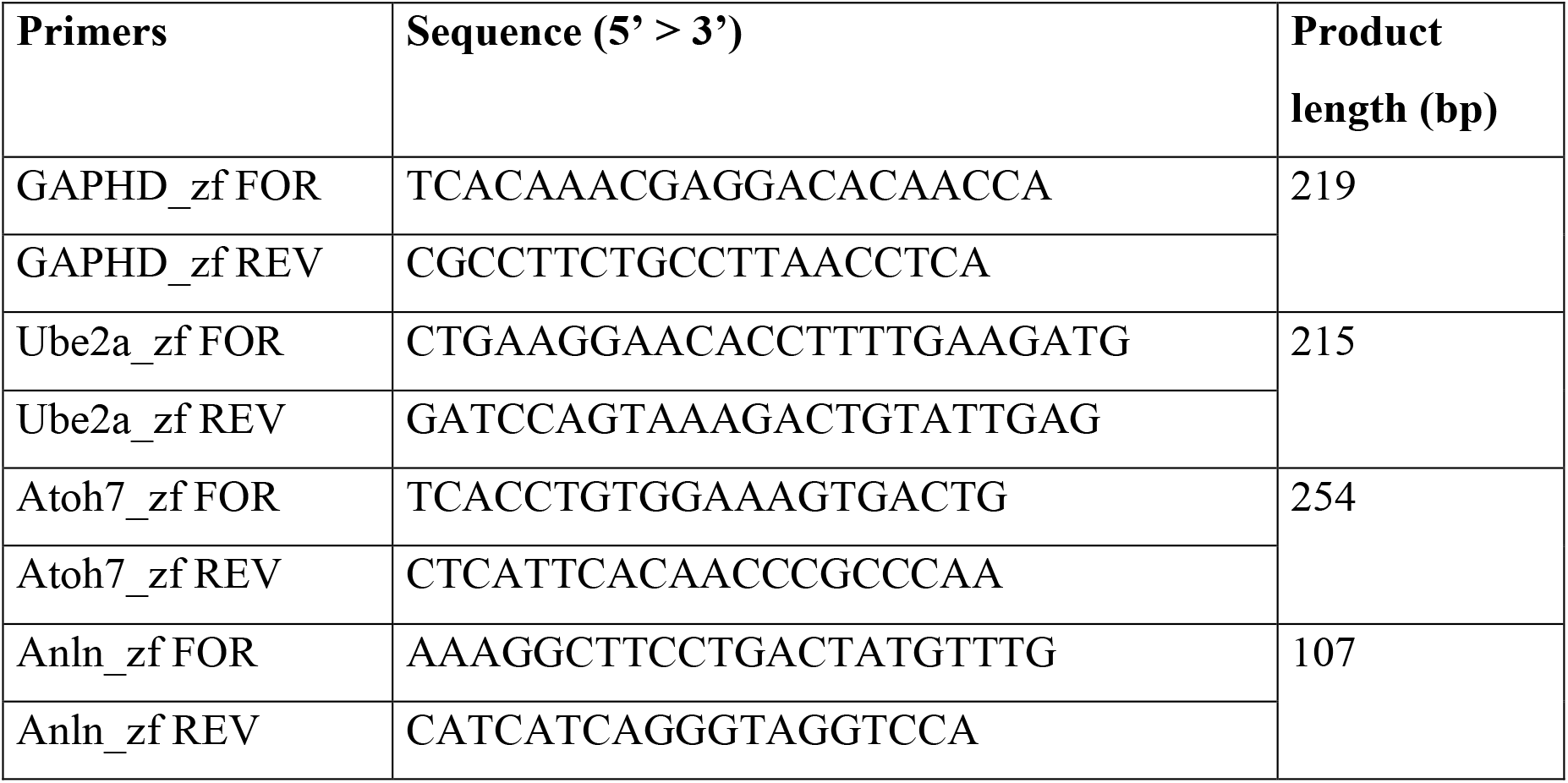
List of primers with amplicon sizes used for quantitative Real-Time-PCR.

### Functional category enrichment analysis and network analysis

Functional enrichment analysis was performed with Metascape ^80^ using the Danio rerio EnsEMBL IDs of the list of differentially regulated genes as “Input as species” and “Analysis as species” species through the custom analysis mode. Enrichment analysis was performed against GO “Biological Process” using P Value cutoff of 0.05 and otherwise default parameters. Human disease annotation was performed with Metascape using *Danio rerio* as “Input as species” and *H. sapiens* as “Analysis as species”. Under the “Annotation” mode, all repositories under Genotype/Phenotype/Disease were selected for disease annotation. To rule out potential sampling or biological bias, we performed an additional enrichment analysis by restricting the background to the list of expressed genes as detected by the arrays, using both Metascape and KOBAS ^81^. Network interactions between the differentially expressed genes were retrieved through the STRING database ^82^, “multiple proteins” mode and default parameters otherwise. Network interactions were visualised using Cytoscape ^83^

### Weighted Gene Co-expression Network Analysis

The pipeline proposed by Langfelder and collaborators ^84^ in their CRAN package was followed to infer gene co-expression networks and identify network modules within R 3.6.3 statistical environment. Networks were inferred using the TOMsimilarityFromExpr function with “cor” as gene coexpression measure. The soft-threshold parameter was optimized with the function pickSoftThreshold and the best threshold (α=16) selected by visual inspection in order to follow a scale-free topology model, as suggested by the WGCNA pipeline. Correlations between modules eigengenes, status and library batch were computed. Modules with the highest correlation for status and significant p-value (α<=0.05) were selected for further analysis. Within the selected modules, highly connected structure of submodules were identified using the leading eigenvector community detection method ^85^ implemented in the igraph package (v1.2.5) for R (http://igraph.com).

### Whole mount Immunohistochemistry

For immunohistochemical labeling, embryos were fixed in 4 % PFA for 1 hour at room temperature or overnight at 4°C. Embryos were washed 3 times in PTw and kept for a week maximum in PTw. Fixed embryos were blocked in blocking solution (10 % goat serum, 1 % bovine serum albumin, 0.2 % Triton X-100 in PBS) for one hour. Embryos were permeabilized with 0.25 % trypsin-EDTA (1X, Phenol red, Gibco – Life Technologies) on ice for 5 min. Primary (mouse anti-β-catenin, 1:100, BD Biosciences Cat. No 610153) and secondary (anti-mouse IgG conjugated to Alexa Fluor 488, 1:250, Invitrogen) antibodies were added for 2 overnights each and DAPI was added from the first day of incubation in the antibody mix. Stained embryos were kept in PTw at 4°C in dark until imaging. Embryos were embedded onto a 35 mm Glass-bottom Microwell dish (p35G-1.5-10-C, MatTek) and oriented with a femtoloader tip (eppendorf) in the position needed for imaging until the agarose had polymerized. Confocal imaging was performed using a laser scanning confocal microscope Leica SpE using a Leica 40X, 1.15 NA oil-immersion objective.

For the analysis of the intensity along the apical-to-basal membrane, a line of a defined length along the apical-to-basal membrane was drawn for 9 cells at 3 different z-sections for each embryo. Signal intensities were obtained using Fiji and average values for the 9 cells were calculated. For the analysis of the intensity along the apical membrane, the number of peaks were counted after Ctnnb1 signal intensity measurement. For the signal intensity measurements, a line of defined length was drawn along the apical membrane of the retina on Fiji and signal intensities were retrieved. The length of line was the same for all z-sections of an individual embryo. Normalization was performed by the highest value for each line. Measurements were obtained on 3 different z sections for each embryo.

## RESULTS

### Transcriptome analysis of wild type and *lakritz* identified 137 statistically significant differentially expressed genes

Expression of *atoh7* in the retina is first detected at around 25-28 hpf and it reaches its peak at around 36 hpf ^86^. The earliest post-mitotic RGCs in the retina are detected at around 28 hpf, a developmental time-point corresponding to the earliest onset of retinal differentiation ^87^. To identify early Atoh7-regulated genes, transcriptome analysis was performed on eyes from single *lak*^*−/−*^ mutant (*lakritz*) ^6^ and wild type zebrafish embryos at 28-30 hours post fertilization (hpf) based on Affymetrix microarrays (see methods and **Fig. 1**).

Differential gene expression analysis performed with LIMMA resulted, after batch effect removal, with 171 significantly differentially expressed probes (adj. P-value < 0.05) (**Fig. 2A**) corresponding to 137 genes annotated onto EnsEMBL database (**Fig. 2B and Supplemental Table 1**). Among these, we confirmed down-regulation of *atoh7* in the *lakritz*, consistent with its known role as self-activator ^33^. Also consistent with the presence of *bona fide* Atoh7-regulated targets in the 137 cohort is the presence of additional 7 genes, which have been previously reported to contain a well-characterised Ath5 consensus binding site ^33,53^. Among the 8 Atoh7-direct targets, besides *atoh7* itself, the thyrotroph embryonic factor *tefa* ^88^ and the atypical cadherin receptor 1 *celsr1a (CELSR1)* ^89^ were down-regulated in the *lakritz*, suggesting their positive regulation by Atoh7. Conversely, the retina and anterior neural fold homeobox transcription factor *rx1 (RAX)* ^90^ the Wnt signalling pathway regulator *notum* ^91^ the transmembrane protein *tmem165* ^92^ and the F-actin binding protein and cytokinesis regulator *anillin (ANLN)* ^93^, were upregulated in the *lakritz*, suggesting that they are negatively modulated by Atoh7.

**Figure 2:**
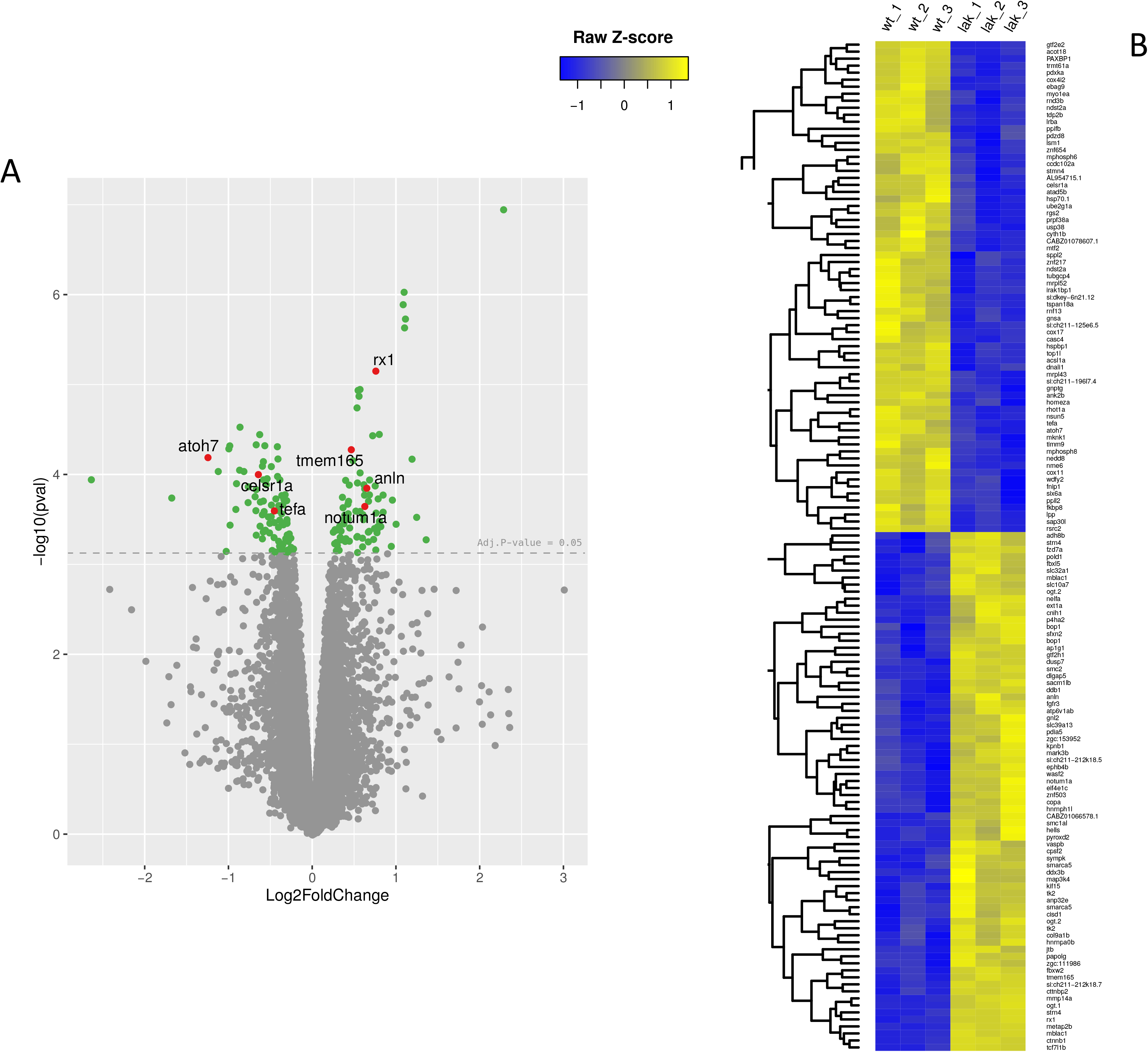
Volcano and heatmap of differentially expressed genes in lakritz vs wild-type eyes. *A)* Volcano plot highlighting Atoh7 and its direct targets (in red) among all differentially expressed probes (in green) with adjusted p-value < 0.05. *B*) Heatmap was constructed by calculating row Z-score using normalised log2 intensities of 144 of the 173 differentially expressed probes with corresponding gene annotation, using complete hierarchical clustering in R.

### Functional category enrichment reveals neural retina, cell cycle and Wnt pathways regulated downstream of Atoh7

Functional enrichment analysis in Metascape using default parameters reveals “neural retina development” (GO:0003407) as the most highly significantly enriched GO Biological Process category, consistent with the role of Atoh7 as regulator of retinal development **(Fig. 3A and Supplemental Table 2)**. To further increase stringency, ruling out sampling or biological bias in the analysis, the background was restricted to the list of expressed genes as detected by the arrays ^81^ both using Metascape and Kobas 3.0, which incorporates knowledge from 5 pathway databases (KEGG PATHWAY, PID, BioCyc, Reactome and Panther), and 5 human disease databases (OMIM, KEGG DISEASE, FunDO, GAD and NHGRI GWAS Catalog). This analysis consistently underscored “neural retina development” (GO:0003407) as the enriched term both by Metascape (not shown) and Kobas (**Supplemental Table 3**). This category contains 9 statistically significantly differentially expressed genes (including *atoh7*), which comprehend early expressed eye-field transcription factors (EFTFs, *rx1* and *six6a*) ^94–97^, stress response and extracellular matrix remodelling factors (*hsp70.1*, *mmp14a*) ^98–101^, chromatin regulators (*smarca5*) ^102^ as well as microtubules organisers and cell cycle regulatory proteins (*tubgcp4*, *znf503*, *gnl2*) ^103–105^. Other (albeit less significant) relevant biological processes emerging from this analysis were “cell cycle process” (GO:0022402), “chromatin remodeling” (GO:0006338) and “Wnt signaling pathway” (GO:0016055) (**Fig. 3B and Supplemental Table 2)**.

**Figure 3:**
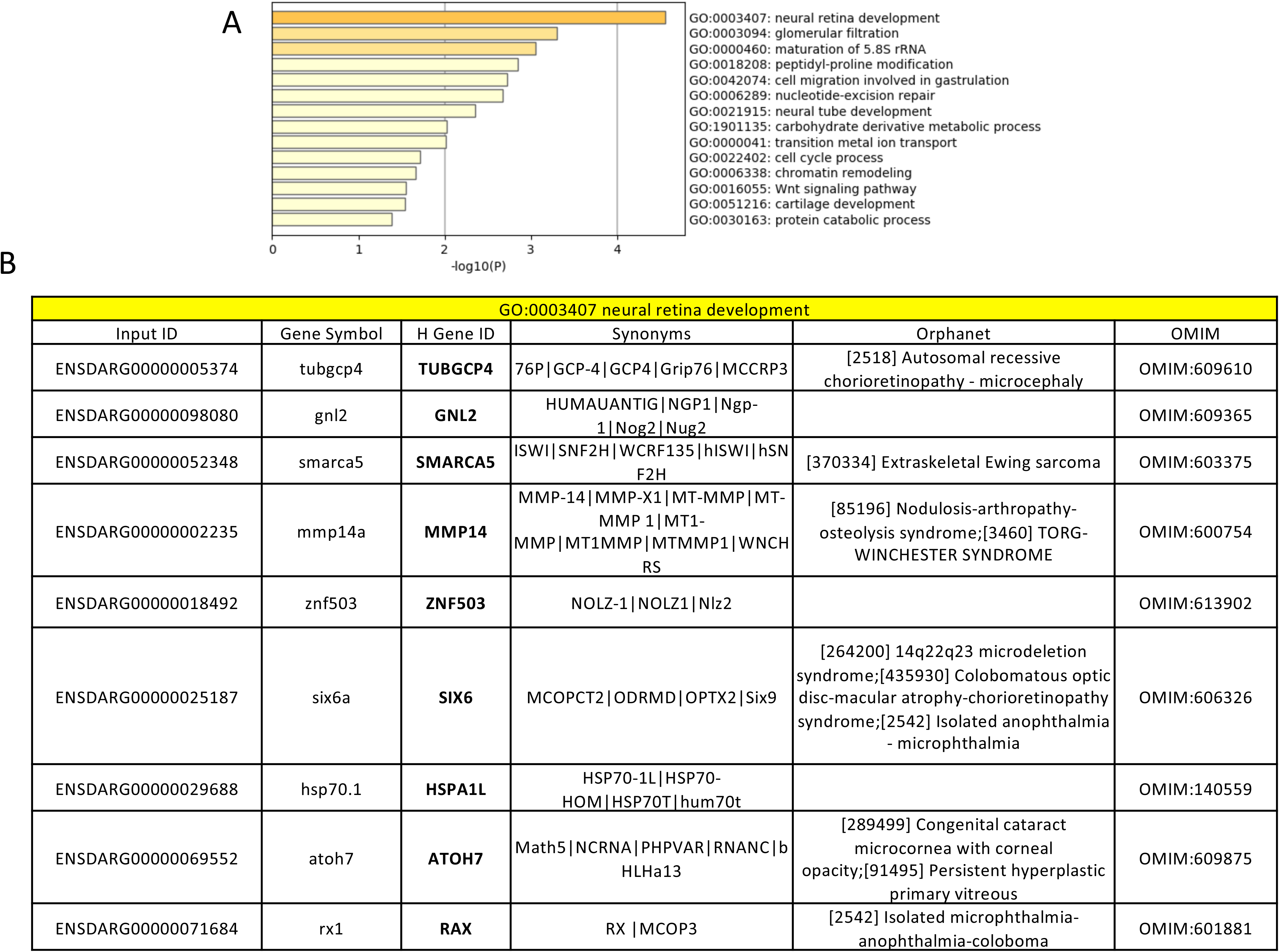
Functional enrichment analysis. *A)* Statistically significantly overrepresented GO Biological Process categories (Metascape). *B)* Significantly differentially expressed genes belonging to the “neural retina development” category **(see also Supplementary Table 2)**.

We next investigated the known relationships amongst the 137 Atoh7-regulated genes via network reconstruction analysis (see methods). Analysis conducted with STRING-DB v 11.0 s highlighted three main gene subnetworks, which are suggestive of the over-represented GO Biological processes, namely “retinal development”, “cell cycle” and “Wnt signalling pathway” **(Fig. 4)**. Furthermore, this analysis suggested *atoh7, rx1 and six6a* as the core of a retinal “kernel” composed of early developmental eye-specific transcription factors ^106^ **(Fig. 4)**.

**Figure 4:**
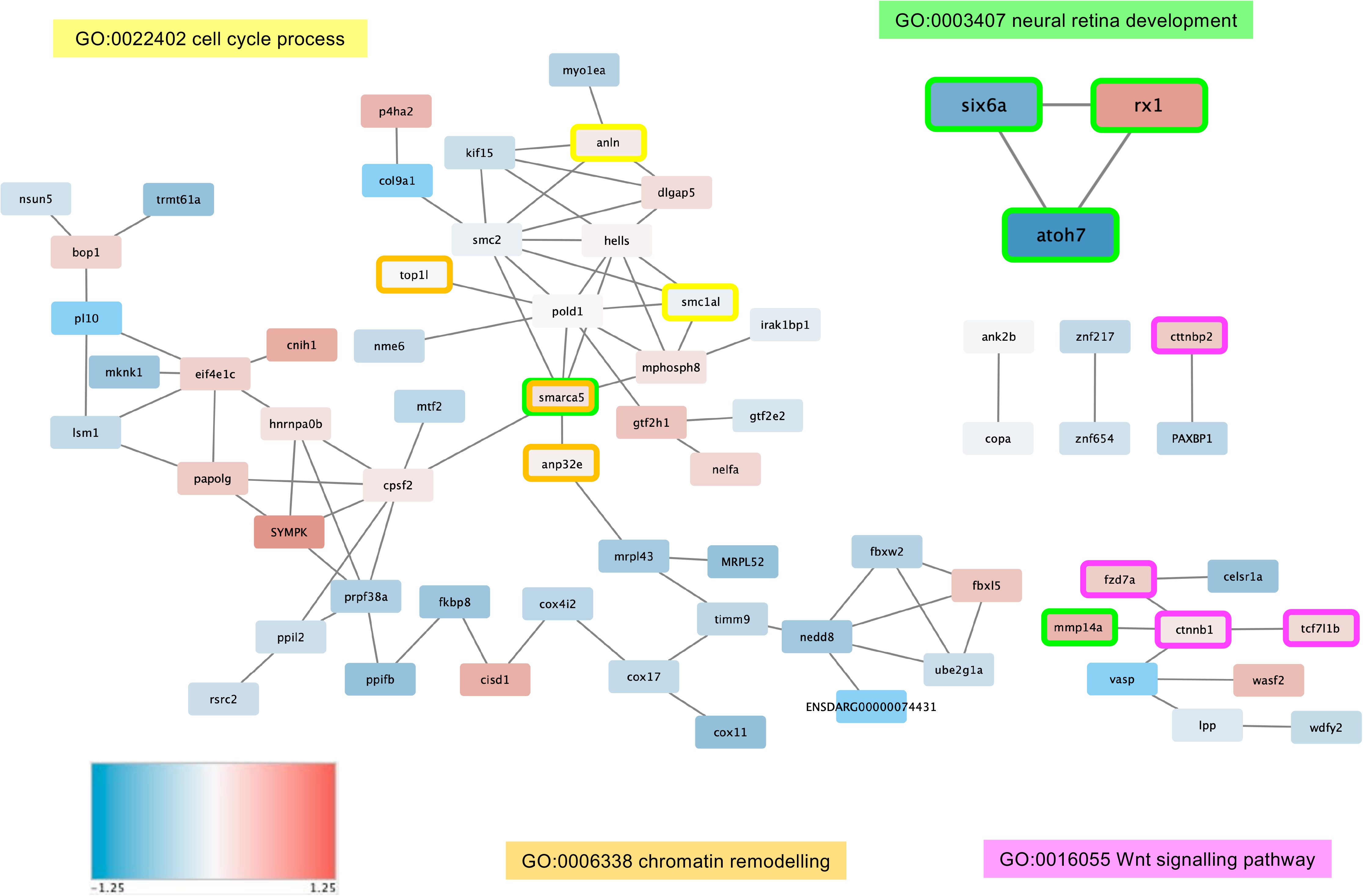
Interaction network downstream of Atoh7. Known interactions between downstream targets of Atoh7 from the STRING database and visualised with Cytoscape (genes without known interactions are not represented). Node colours represent the log2 fold-change of the gene expression in *lakritz* versus wild type eyes. Node borders are coloured by gene ontology annotation: “neural retina development” (green), “cell cycle process” (yellow), “Wnt signalling pathway” (pink) and “chromatin remodelling” (orange).

### Weighted Gene Co-expression Network Analysis revealed a coexpression module with a cluster of genes tightly interconnected to Atoh7

To explore global changes of gene expression in the *lakritz* mutant, we conducetd a weight gene co-expression network analysis (WGCNA) ^84^ to the available microarray data. Notwithstanding the small number of samples, we identified 16 recurrent functional modules based on co-expression pattern analysis on the full transcriptome dataset. In order to identify co-expression modules significantly affected by the *lakritz* mutation, we tested for their association with available covariates, including batch and mutation status. Out of the 16 modules found, two of them showed a high correlation with the mutation condition (wild type vs *lakritz*). Specifically, modules 13 (overall up regulated) and module 3 (overall down regulated) were found highly significant – with a correlation of 0.97 (p=0.002) and −0.99 (p=1e-5), respectively (**Supp. Fig. 2 and Supplemental Table 4**). Functional network analysis by STRING DB (https://string-db.org/cgi/network.pl?taskId=hNkdhEXQnBhR) on M13 (the smallest module, which we also found to contain *atoh7*) revealed an enrichment in “eye morphogenesis (blue color in **Supp. Fig. 3**), “retina layer formation” (red color in **Supp. Fig. 3**), and a cluster of genes previously found to be dysregulated in “light responsive, circadian rhythm processes” (PMID: 20830285 and PMID: 21390203) (light and dark green color in **Supp. Fig. 3**).

Lastly, we analyzed in detail the topology of the Module 13. We identified 4 submodules with high within-community connectivity, which show decreasing degree of connectivity from left to right **(Fig. 5).** The submodule containing *atoh7* (left) shows a densely interconnected cluster of genes with high topological overlap. These genes likely participate in common regulatory and signaling circuits including retina layer formation (e.g. *atoh7, rx1*) and Wnt/β-catenin signalling pathway (e.g. *fxd7a, tcf7l1b*).

**Figure 5:**
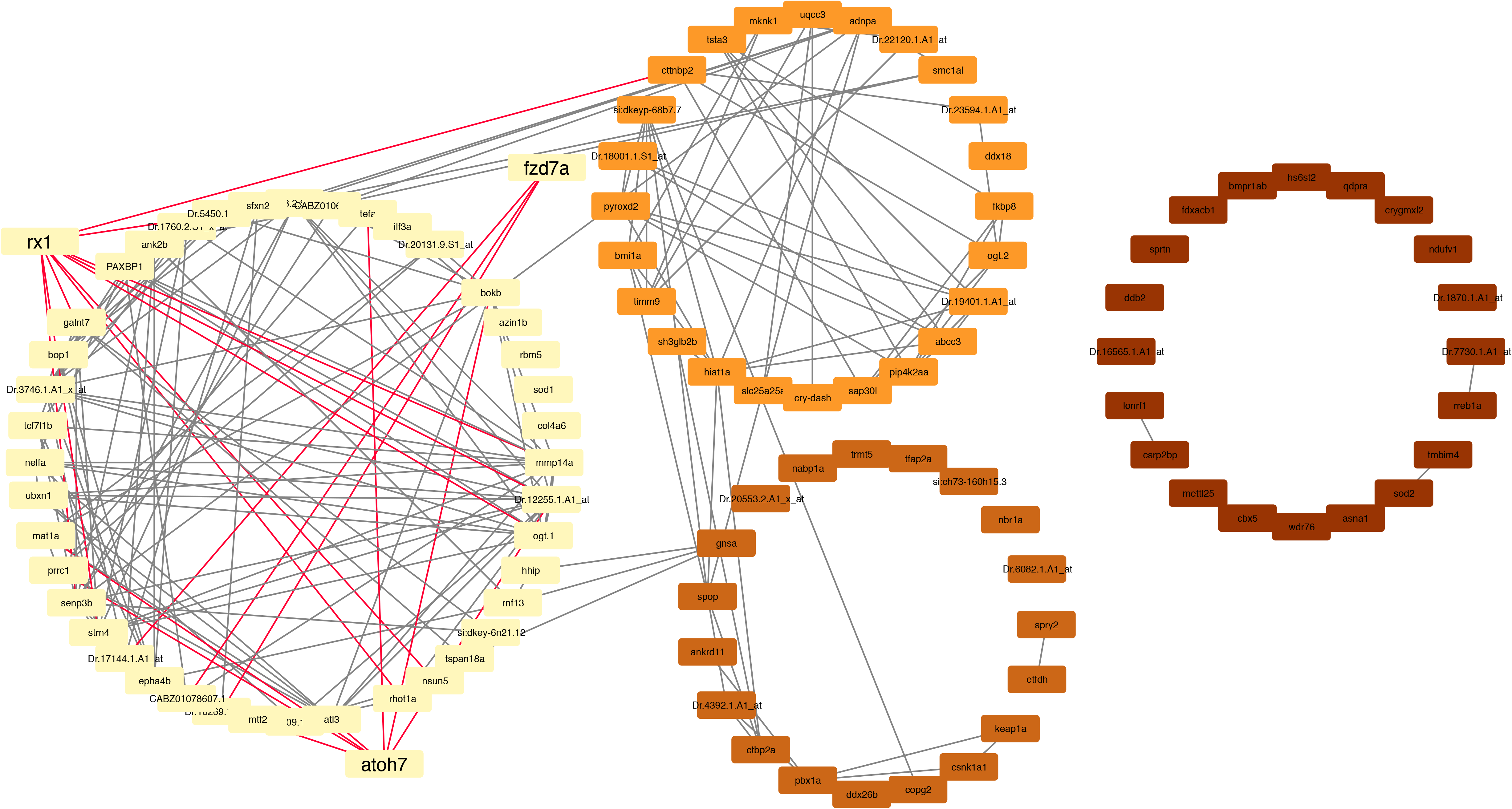
Detailed topology of Module 13. Each node represents a gene while a connection represents a co-expression between two genes (only the first 200 edges in order of co-expression weight were retained for visualization purposes). Submodules are shown with decreasing degree of connectivity from left to right. Highlighted edges represent the connection between retina layer formation and wnt - related genes.

## DISCUSSION

Our differential gene expression analysis of transcriptome data revealed 137 genes that are significantly differentially expressed between *lakritz* and wild type eyes from embryos at a developmental time-point corresponding to the onset of RGC differentiation. We also applied multiple bioinformatics pipelines to perform a functional classification and network reconstruction of the differentially expressed genes. Notably, all methods here applied consistently highlighted “neural retina development” (GO:0003407) as the most biological pathway differentially affected by the *lakritz* mutation. Likewise, the interplay *atoh7*, *rx1* end *six6a* – early developmentally regulated EFTs – consistently emerged as the “kernel” network of this cluster.

The homebox transcription factor Rx1 is well known for its evolutionarily conserved role in the generation and maintenance of multipotent RPCs during morphogenesis and differentiation of the vertebrate eye ^95,97,107–110^. Mutant variants of *RAX* family genes have been linked to congenital developmental eye disorders, particularly microphthalmia, further confirming an early requirement for RPC proliferation and stemness ^29,111–113^. Based on these findings, the data from our analysis suggest that balancing RPCs competence and RGC fate commitment requires *rx1* downregulation by Atoh7. Notably, Atoh7-mediated downregulation of *rx1* is likely direct, since previous *in silico* analyses highlighted the presence of an Atoh7-binding motive in the *rx1* gene cis-regulatory regions ^33^.

Conversely, *six6* genes have been reported as RPCs competence factors, on the one hand suppressing stemness and proliferation via Wnt/β-catenin signalling downregulation, on the other hand promoting expression of RGC differentiation genes ^94^. These findings are in agreement with the present study reporting *six6a* being an Atoh7-upregulated gene. Remarkably variant forms of *SIX6 (SIX9/OPTX2)* have been linked to congenital microphthalmia as well as to the development of glaucoma ^20,29,32,114–119^. Given that mutations in *ATOH7* have been also associated with similar global eye disorders ^14,120^, these observations strongly point at the importance to further understand the interplay *ATOH7*, *RX1* and *SIX6* during eye development, and to assess how disruption of this evolutionarily conserved genetic network might be linked to such eye disorders ^121^.

Besides *atoh7*, *rx1* and *six6a,* significant differentially expressed genes annotated with “neural retina development” were *tubgcp4*, *gnl2*, *smarca5*, *mmp14a*, *znf503*, and *hsp70.1*. The Atoh7-upregulated gamma-tubulin complex protein 4 encoding gene *tubgcp4* is of great interest. Variants of *TUBGCP4* have been linked to autosomal-recessive microcephaly with chorioretinopathy, which comprise a spectrum of eye developmental anomalies including microphthalmia, optic nerve hypoplasia, retinal folds, and absence of retinal vasculature ^103^. This raises the possibility that regulation of this gene might link Atoh7 to retinal-vascular development as well as retinal neurogenesis. Genetic evidences for the GTPase and zinc finger transcriptional repressor encoding genes, *gnl2* and *znf503 (NOLZ1)* as eye disorders-related genes are still missing but studies support their functional requirement for retinal developmental processes, including proper cell cycle exit of RPCs during RGC differentiation ^104,105^. Notably, *tubgcp4* and *gnl2* and *znf503* were implicated in the regulation of cytokinesis late in mitosis ^103–105^. This observation further indicates Atoh7-requirements for the regulation of cell cycle progression, at least in RPCs. The SWI/SNF chromatin remodelling factors have been reported as crucial regulators of the transition from multipotent to committed progenitor and differentiated cell states in multiple eye tissues, with potential implications for eye disorders ^122–125^. The finding of *smarca5* as an upregulated gene in the *lakritz* indicates the importance to address functional implications of this gene for *atoh7*-related eye disorders. Furthermore, the “neural retina development” Gene Ontology category encompassed the reportedly stress response genes *hsp70.1(HSPA1L)* ^101^ and *mmp14* ^126^ as significantly differentially down- and up-regulated, respectively, by the *lakritz* mutation. The crystallin related, heat shock Hsp70 family proteins are emerging as important regulators of RGC survival and regeneration as well as retinal vascular remodelling factors ^101,127–129^. The intriguing finding that *hsp70.1* is highly enriched among the Atoh7-upregulated genes in the “neural retina development” gene cluster supports the idea that upregulation of these stress-response proteins might be relevant also during RGC development. Lastly, the extracellular matrix remodelling factor Mmp14 is reportedly a crucial regulator of cell stemness and vascular remodelling ^130^. Studies have also implicated Mmp14a in retinal developmental processes such as RGC axon guidance and innervation of the optic tectum ^98–100^. Future studies will assess the functional implication of Atoh7-mediated downregulation of *mmp14a* during vascular-retinal development. Interestingly, our network analysis reveals that *mmp14a* is linked with known components of the Wnt signalling pathway, further underscoring the importance of the interplay of this pathway and Atoh7.

The “Wnt signalling pathway” is the second Atoh7-dependent subnetwork emerging in our functional network analysis; which is centered around the *ctnnb1* (β-catenin) gene **(Fig. 4).** Studies have shown that the Wnt/β-catenin pathway promotes RPC proliferation and stemness whilst suppressing *atoh7* activation and RGC differentiation ^131–133^. We here find that the main differentially expressed components of this pathway, namely *ctnnb1*, *fxd7a* and *tcf7l1b*, are upregulated in the *lakritz* mutant, suggesting Atoh7 requirement in the Wnt/β-catenin pathway downregulation **(Fig. 4)**. Concordantly, studies have reported that *atoh7*-expressing retinal progenitors contain low levels of expression in Wnt/β-catenin pathway components when compared with non-*atoh7*-expressing progenitor cells ^35^ further suggestive of a negative feedback regulatory loop integrating Atoh7 and Wnt/β-catenin signalling. Conversely, the planar cell polarity (PCP) signalling component *celsr1a/flamingo*, which has been reported as key regulator of neuronal cell differentiation, neurite outgrowth and axon guidance ^134^ emerges as an Atoh7-upregulated gene in our cohort. A number of studies reported dysregulation of Wnt signalling being associated with retinal diseases ^133,135–137^. Likewise, Wnt, Fzd7/β-catenin pathway has been reported as an important modulator of retinal vascular remodelling ^138^. Further research will clarify the genetic regulatory networks integrating Atoh7 and Wnt/β-catenin signaling in controlling multiple eye tissue development in the vertebrate, and how their dysregulation might results in multiple ocular disorders ^139^.

Previous studies highlighted the importance of the interplay of Atoh7 with components of the Notch signalling pathway for RGC development and regeneration ^43,50,140^. In particular, downregulation of Notch signalling pathway has been proposed as a general mechanism whereby RGC genesis can be enhanced ^140–143^. According to these findings, Notch pathway components (*hes6*, *mib1*, *hdac*, *adam17b*, and *Notch1a*) appear affected by the *lakritz* mutation in our gene expression microarray data **(Supplementary Table 1)** but their expression does not change significantly between wild type and *lakritz* condition in our analysis **(Supplementary Table 1)**. However, even though they fail short of the significance threshold of adjusted p-value < 0.05, it is worth noting that their values suggest a trend in downregulation of Notch pathway-related genes **(Supplementary Table 1).** We suppose that such lack of significance in the differential expression might be linked to the developmental stage used for this analysis. Likewise, the extracted cohort of significantly regulated genes does not comprehend some of the reported RGC maturation-associated factors, such as Cxcr4b, Elavl3 and Isl1 ^33,35,37,53^. Nonetheless, for *cxcr4b* and *elavl3a*, −1.43 FC (nominal p-value of 0.0018) and −1.39 FC (nominal p-value 0.0066) was observed, respectively, consistently with a positive regulation by Atoh7 (downregulated in the *lakritz*). We therefore suppose that expression of such RGC maturation-related factors downstream of Atoh7 might be too low at the developmental time-point selected for this study, to be detected within the chosen significance range.

In support of the Atoh7-dependent transcriptional regulation of progenitor cell division and developmental progression, a third Atoh7-dependent subnetwork emerged in our functional network analyses, which includes cell-cycle and chromatin regulators **(Fig. 4)**. One *bona fide* gene of great interest in this subnetwork is the F-actin binding and cytokinesis regulator Anillin (*ANLN*) ^53,144,145^. Anillin has attracted increasing attention as a potential disease-related gene (ORPH:93213, OMIM:616027). Evidence point at *anillin* expression levels being associated with cell proliferation and migration disorders in cancer and kidney diseases ^146–149^. Additional roles for this actin binding protein have been reported in nerve cell development ^150,151^ and dysregulation of Anillin has been implicated in central nervous system myelin disorders ^152,153^. In the zebrafish retina, *anillin* expression levels appears required to favour cell cycle progression and restrict RGC genesis in Atoh7-expressing RPCs ^53^. Concordantly, in the presence of *anillin* downregulation many more RPCs turn on *atoh7* and become RGCs ^53^. We here find *anillin* as an Atoh7-downregulated gene, suggesting that a molecular feedback regulatory loop of an as yet unknown nature between *anillin* and *atoh7* is at work, to balance RPCs developmental progression. Whilst further investigations will address this question, the functional network analyses from this study suggest that the Wnt/β-catenin signalling pathway might be involved. Studies have shown that *anillin* expression is positively associated with the expression of β-catenin (*ctnnb1*) ^154^ consistent with *ctnnb1* also being an Atoh7-downregulated gene in our gene cohort. Furthermore, Anillin appears to be an essential regulator of epithelial cell-cell adherens junctions via regulation of the E-Cadherin/β-catenin complex ^148,155,156^. In agreement with this idea, we here make the preliminary observation that *anillin* knock down results in accumulation and displacement of β-catenin signal in the apical and apical-lateral membrane of RPCs **(Supp. Fig. 4A,B)**. Notably, in addition to promoting cell-to-cell adhesion ^157,158^ β-catenin functions as nuclear transcriptional co-activator of Wnt signalling responsive genes promoting proliferation and inhibiting differentiation ^159–161^. Thus, regulation of β-catenin accumulation and localization to the E-Cadherin/β-catenin complex might be a mechanism whereby Anillin controls not only cell adhesion dynamics, but also Wnt signalling pathway activity ^157^. It is also intriguing to note that high levels of *anillin* expression where found in human choroidal endothelial cells, leading to the obvious hypothesis that Anillin is a potential regulator of choroidal angiogenesis ^162,163^. These observations suggest both RPCs and endothelial cell behaviours might require *anillin* expression levels during retinal-vascular developmental interactions. Future studies will assess the functional implications of the interplay of Atoh7, Anillin and Wnt pathway components for the dysregulation of eye developmental processes. In line with this hypothesis is our preliminary observation that *anillin* remains significantly affected (namely upregulated) by the *lakritz* condition, starting from 25 hpf (coinciding with the onset of *atoh7* expression ^86^) until after 72 hpf (when all retinal layers are fully differentiated ^87^). Furthermore, the extent of such upregulation in subsequent developmental stages suggests oscillatory dynamics of *anillin* transcriptional downregulation by Atoh7 **(Supp. Fig. 4C)**. Overall, these observations further underscore the importance of feedback loops involving Atoh7, *anillin* and Wnt signalling, the dysregulation of which could possibly contribute to the development of vascular-retinal disorders.

Finally, we have applied weighted gene co-expression network analysis to explore gene co-expression relationships and identify co-expression modules potentially involved in *atoh7* function. In addition to highlighting the already known interaction networks that were enriched within the 137 differentially regulated genes, we were able to identify two co-expression modules significantly affected by the *lakritz* mutation (**Supplementary Table 4**). One of them in particular (module 13) contains a cluster of highly interconnected genes including *atoh7* itself, *rx1* and members of the Wnt signalling pathway (e.g. *tcf7l1b*, *fzd7a* and *mmp14a)*. This analysis therefore further confirms tight functional interaction between neural retinal development and Wnt pathway genes from our gene differential expression data.

Lastly, our knowledge-based analysis of the M13 members by STRING database further extended these finding by revealing the existence of a network of interactions between *atoh7* and *rx1* dependent “retina layer formation”, “eye morphogenesis” pathways, Wnt signalling pathway components (e.g. *tcf7l1b*, *fzd7a* and *mmp14a)*, and two new gene networks previously found to be dysregulated in light responsive, circadian rhythm processes ^88,164^ **(Supp. Fig. 3)** (https://string-db.org/cgi/network.pl?taskId=hNkdhEXQnBhR). At present, very little is known on the functional importance of circadian clock genes, but increasing evidence indicates their implication in multiple eye tissues developmental processes as well as ocular disorders ^165–169^. This study further supports this evidence, by showing that Atoh7-dependent regulatory networks integrates such circadian clock genes.

In sum, this Atoh7 targets analysis extends data from other studies focussing on transcription factors cascades enhancing RGC differentiation. We here provide new insights on Atoh7-dependent developmental processes that might be regulated in global developmental eye disorders. First, they suggest that Atoh7 directly controls a two-tiered regulatory network balancing early acquisition of progenitor cell competence (*e.g.* through *six6a*, *rx1*) and repression of pro-multipotency and proliferative processes (*e.g.* through chromatin remodelling, cell cycle and Wnt pathway regulation). Secondly, these data highlight for the first time many previously unreported cytoskeletal proteins, chromatin remodelling factors, stress-response proteins, and even circadian clock genes as Atoh7-regulated genes. Third, this analysis underscores both direct and potential functional genetic links of many of these factors to eye developmental disorders. This study thus contributes to laying the groundwork for the identification of key candidate molecules and their networks as potential targets for early eye disease detection and therapeutic applications.

## Supporting information

Complete microarray dataset after normalisation and batch correction

Gene Ontology enrichment output from Metascape (Biological Process)

Weighted gene co-expression network analysis shows Module 3 and 13 with highly interconnected genes.

## Nonstandard Abbreviations

RPCs: retinal progenitor cells
NCRNA: retinal non-attachment
ONH: optic nerve hypoplasia
ONA: optic nerve aplasia
PHPV: persistent hyperplastic primary vitreous
RGCs: retinal ganglion cells
EFTFs: eye field transcription factors

## ACKNOWLEDGEMENTS

We are grateful to W.A. Harris for supporting this study at Cambridge University and Karen Vranizan (University of California) for technical assistance. We also thank I. Pradel and F. Zolessi (University of Cambridge) for technical assistance and F. Zolessi for discussion. We acknowledge S. Sel (University of Heidelberg) for material support. We thank the fish facility management group for fish maintenance and technical assistance. This work was supported by the Wellcome Trust, the Deutsche Forschungsgemeinschaft Research Grant PO 1440/1-1 to L. Poggi, the Landesgraduiertenförderung (Funding program of the State of Baden Württemberg, Germany) to A-L. Duchemin and the Australian Research Council Discovery Project Grants DP140101067 and DP190102771 to M. Ramialison. The Australian Regenerative Medicine Institute is supported by grants from the State Government of Victoria and the Australian Government. M. L. Allende and F. Gajardo were supported by ANID/FONDAP/15090007; F. Gajardo acknowledges the support of CONICYT REDES 150094.

## AUTHOR CONTRIBUTIONS

L. Poggi designed research; G. Covello, A-L Duchemin, F.B. Tremonti, L. Poggi and J.Ngai performed experiments; F.J. Rossello, M. Filosi, F. Gajardo, E. Domenici, M. Eichenlaub, M. Ramialison performed bioinformatic analysis with inputs from J.M. Polo, D. Powell, M. L. Allende; L. Poggi and M. Ramialison wrote the manuscript with inputs from all authors.

---

**Supplementary Figure 1:**
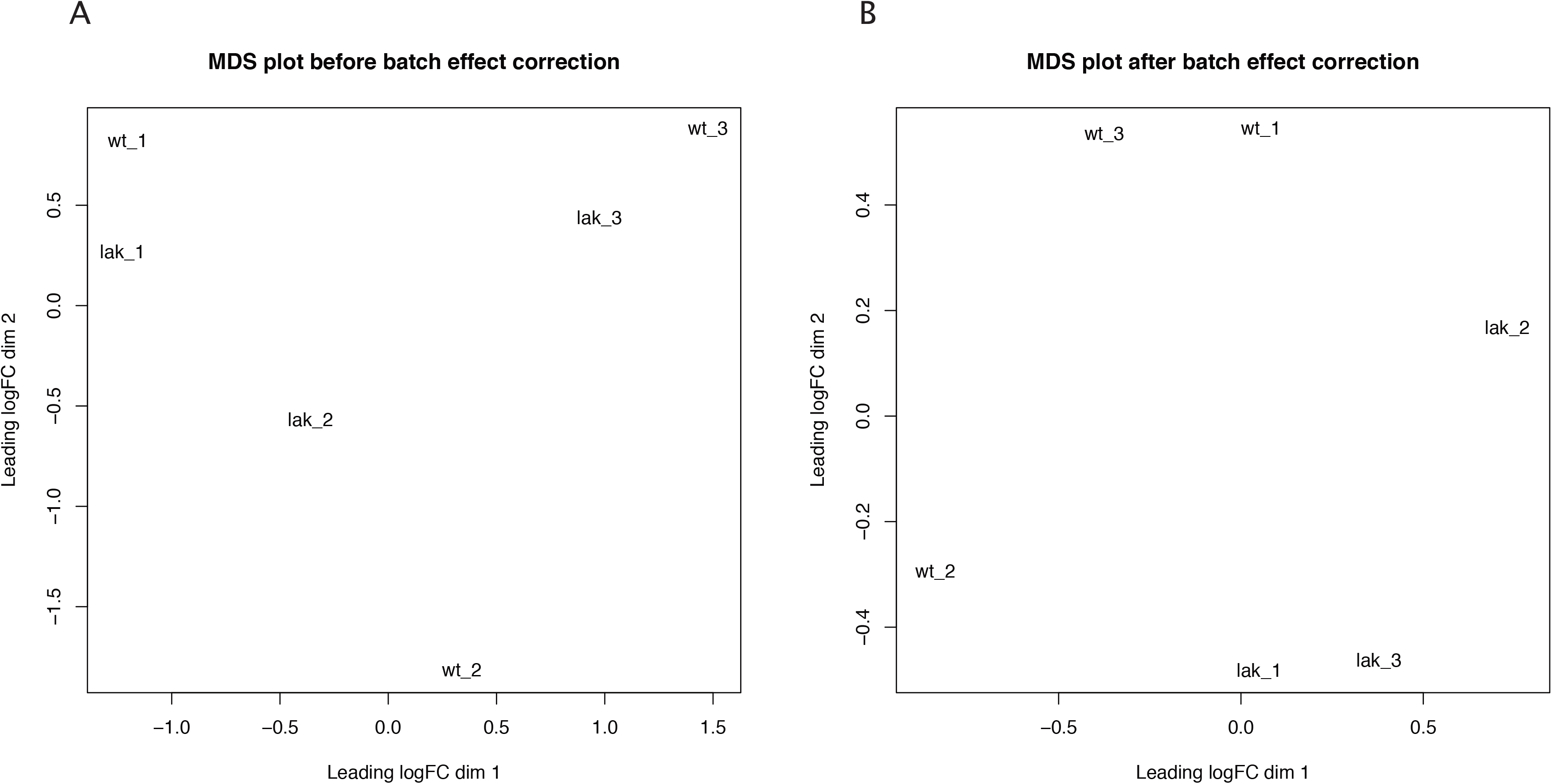
Batch effect correction. MDS plots before *A)* and after *B)* batch effect correction using Combat (plotMDS in R).

**Supplementary Figure 2:**
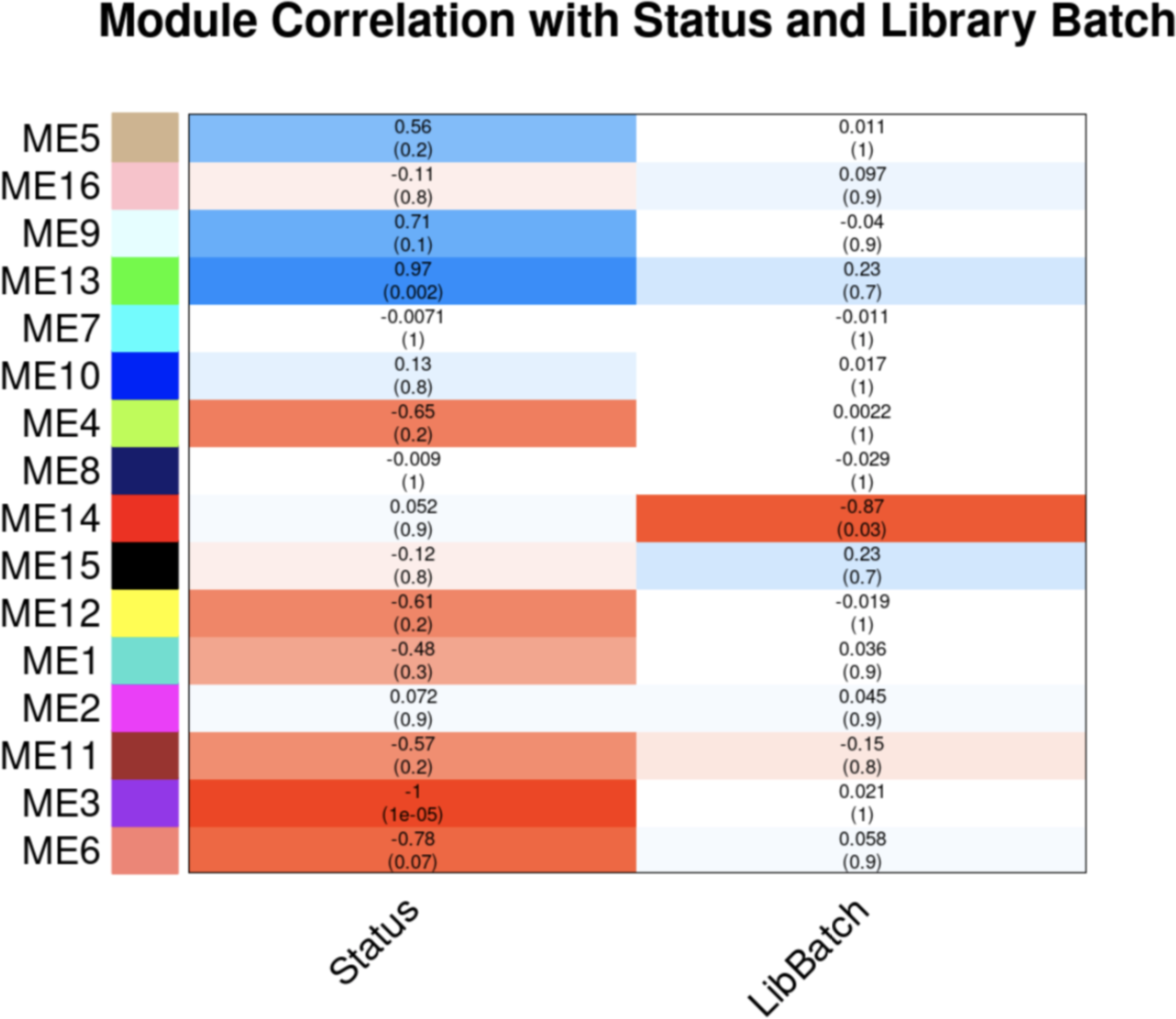
co-expression modules significantly affected by the *lakritz* mutation.

**Supplementary Figure 3:**
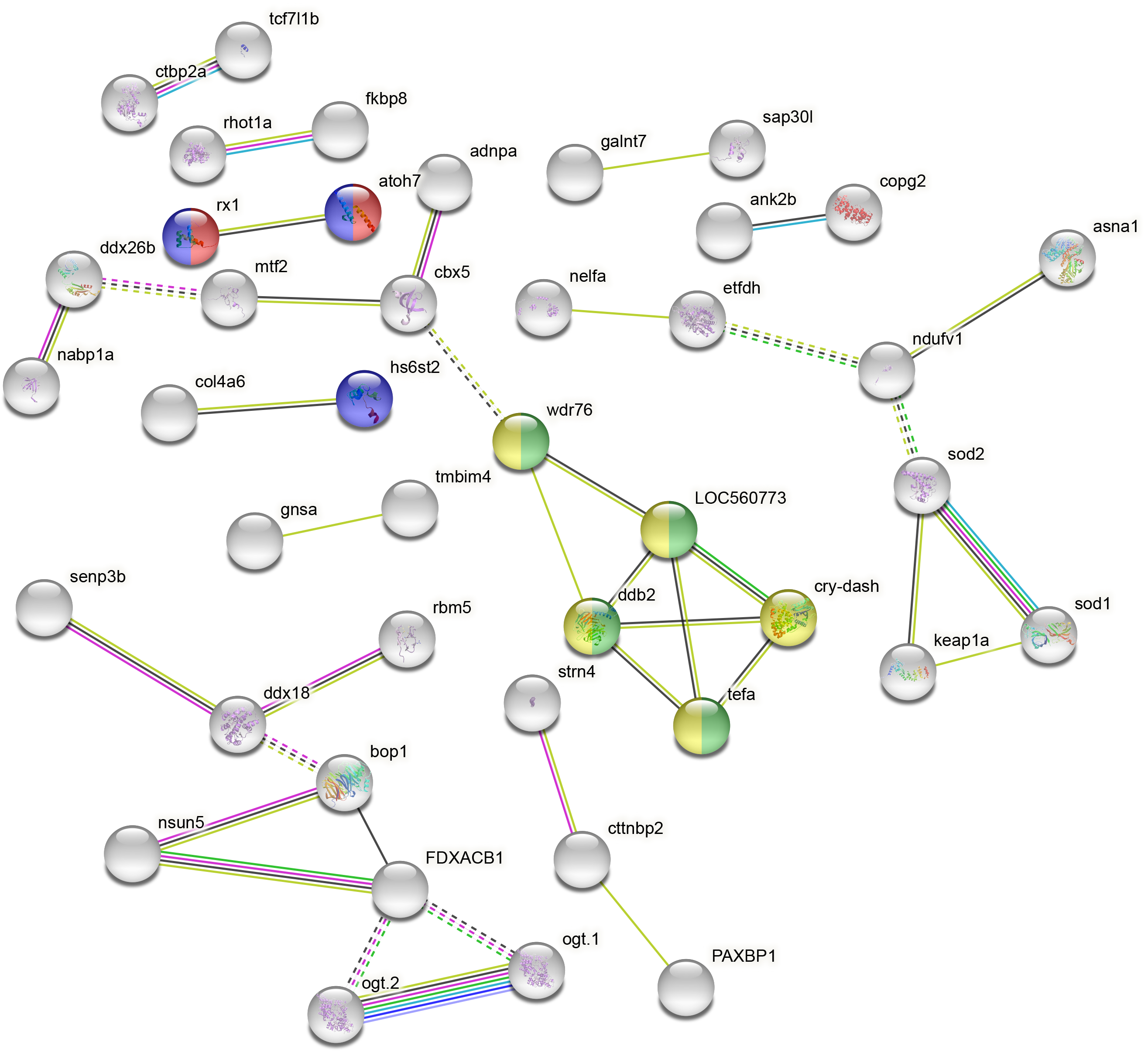
Functional network analysis on M13: (https://string-db.org/cgi/network.pl?taskId=hNkdhEXQnBhR)

**Supplementary Figure 4.**
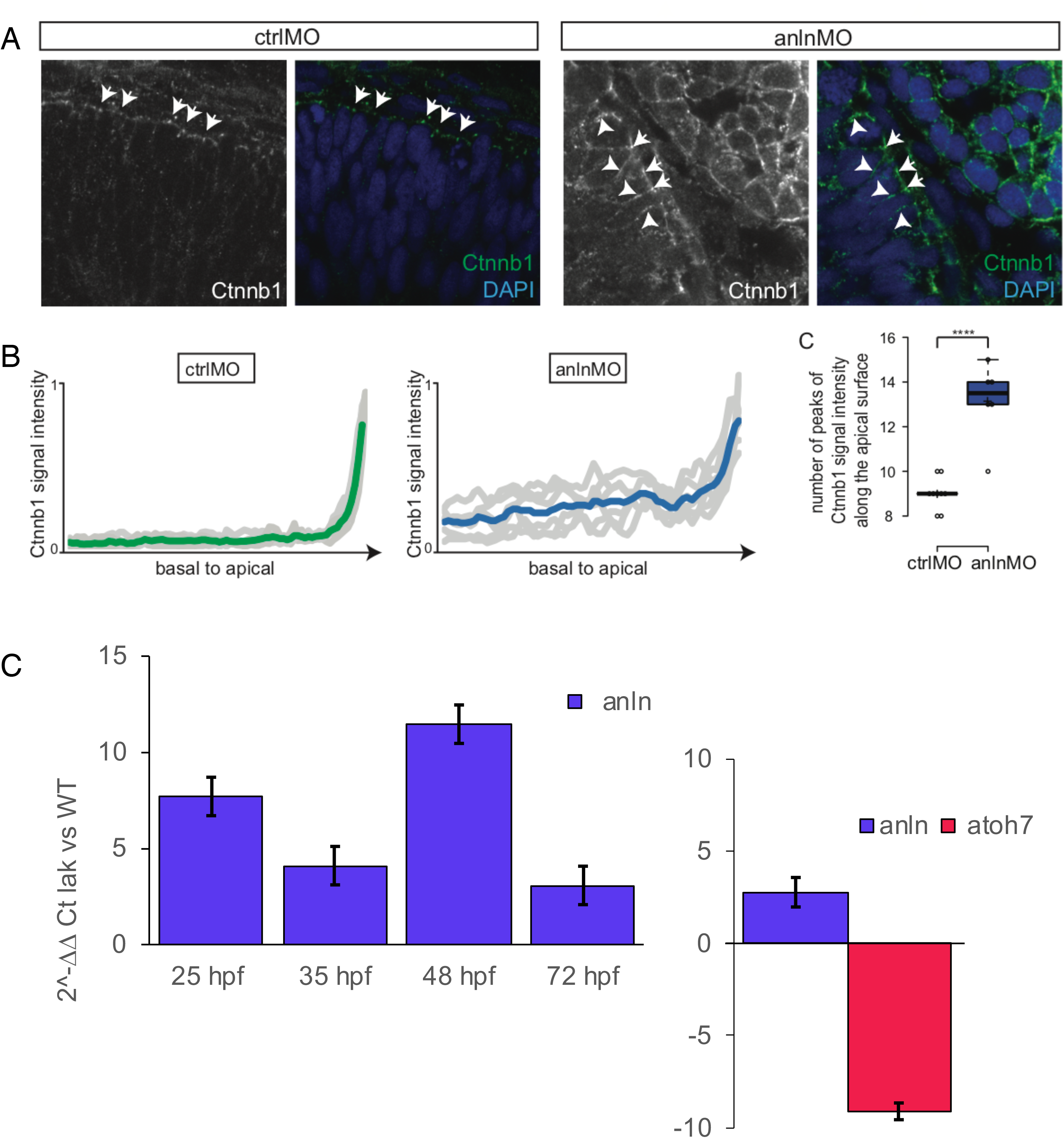
Dysregulation of Ctnnb1 localization by *anilin* knock-down and *anillin* expression dynamics. *A)* Ctnnb1 staining in control (ctrlMO, n=3 embryos) versus anilin knock-down (anlnMO, n=3 embryos) in morpholino injected embryos at 30hpf. Arrows show apical location, arrowheads show apical-to-basal location. *B)* Graph showing the normalized Ctnnb1 intensity signal along the basal-to-apical membrane of the apical-most cells in control (n=3 embryos) versus anlnMO (n=2 embryos) injected embryos. The colored line shows the averaged intensity of all lines for the ctrlMO and anlnMO. Boxplot showing the number of peaks of Ctnnb1 signal intensity along the apical surface in ctrlMO (n=3 embryos) versus anlnMO (n=2 embryos) injected embryos. P<10^−4^. Center lines show the medians; crosses show the means; box limits indicate the 25th and 75th percentiles as determined by R software; whiskers extend 1.5 times the interquartile range from the 25th and 75th percentiles, data points are represented as circles. Student’s t-test. *C) Anillin* mRNA levels show dynamic variations during subsequent developmental stages. qPCR was performed on eyes from *lakritz* or wild type embryos at 25, 35, 48, 72 (left) and 96 hpf (right) to assess the trend of *anillin* and *atoh7* expression. The relative gene expression (*lakritz* vs wild type) was calculated using the CT method for each stage. Histogram values are expressed as mean ± s.e.m. (p < 0.05) and the mRNA levels of both *gapdh* and *ube2a* were used as internal controls. The statistical analysis is described in the methods section.

**Supplementary Table 1: Complete microarray dataset after normalisation and batch correction**.

**Supplementary Table 2: Gene Ontology enrichment output from Metascape (Biological Process)**

**Supplementary Table 3: Kobas-based analysis underscores “neural retina development” (GO:0003407) as the most enriched category.**

**Supplementary Table 4: Weighted gene co-expression network analysis shows Module 3 and 13 with highly interconnected genes.**

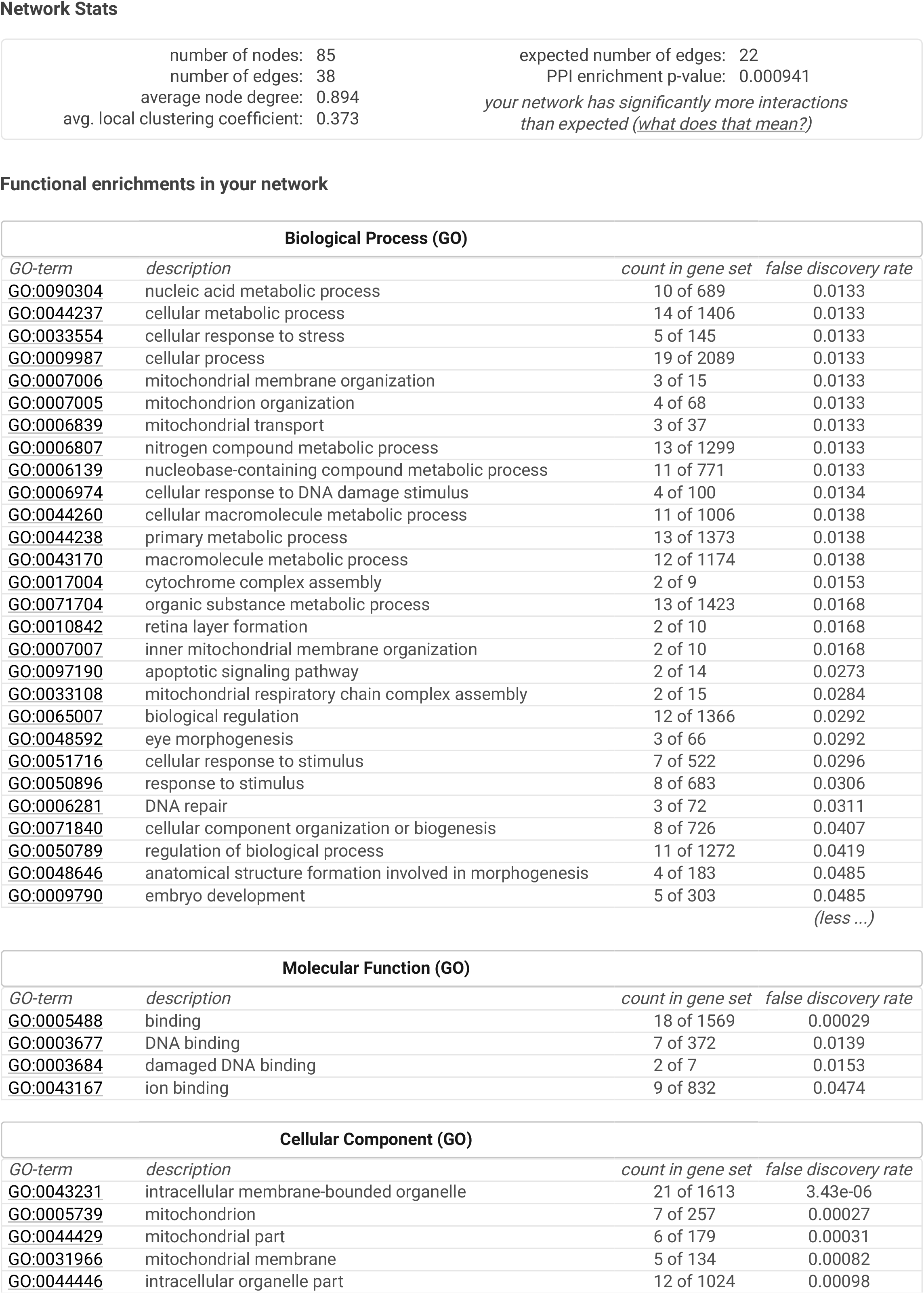

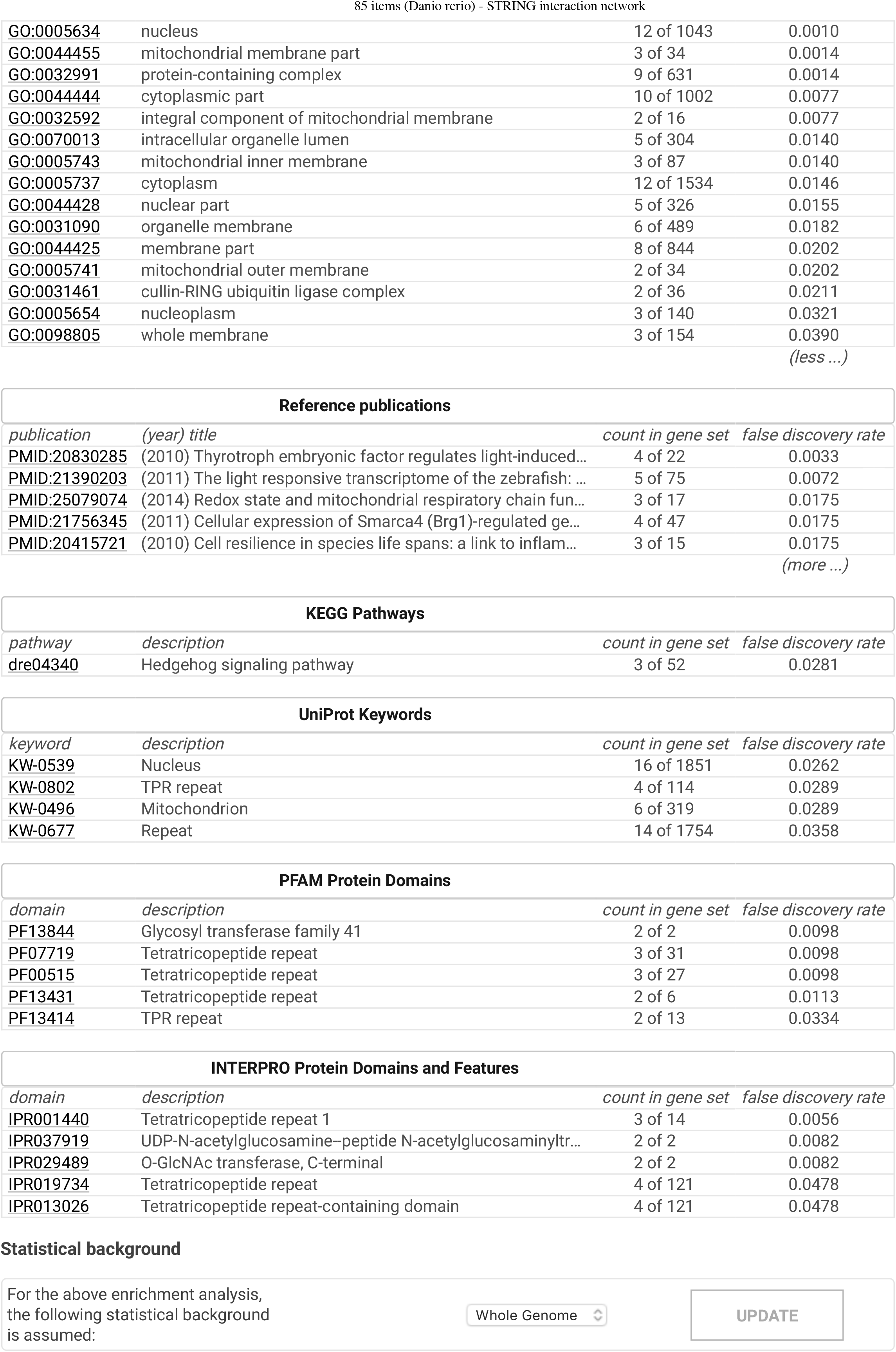

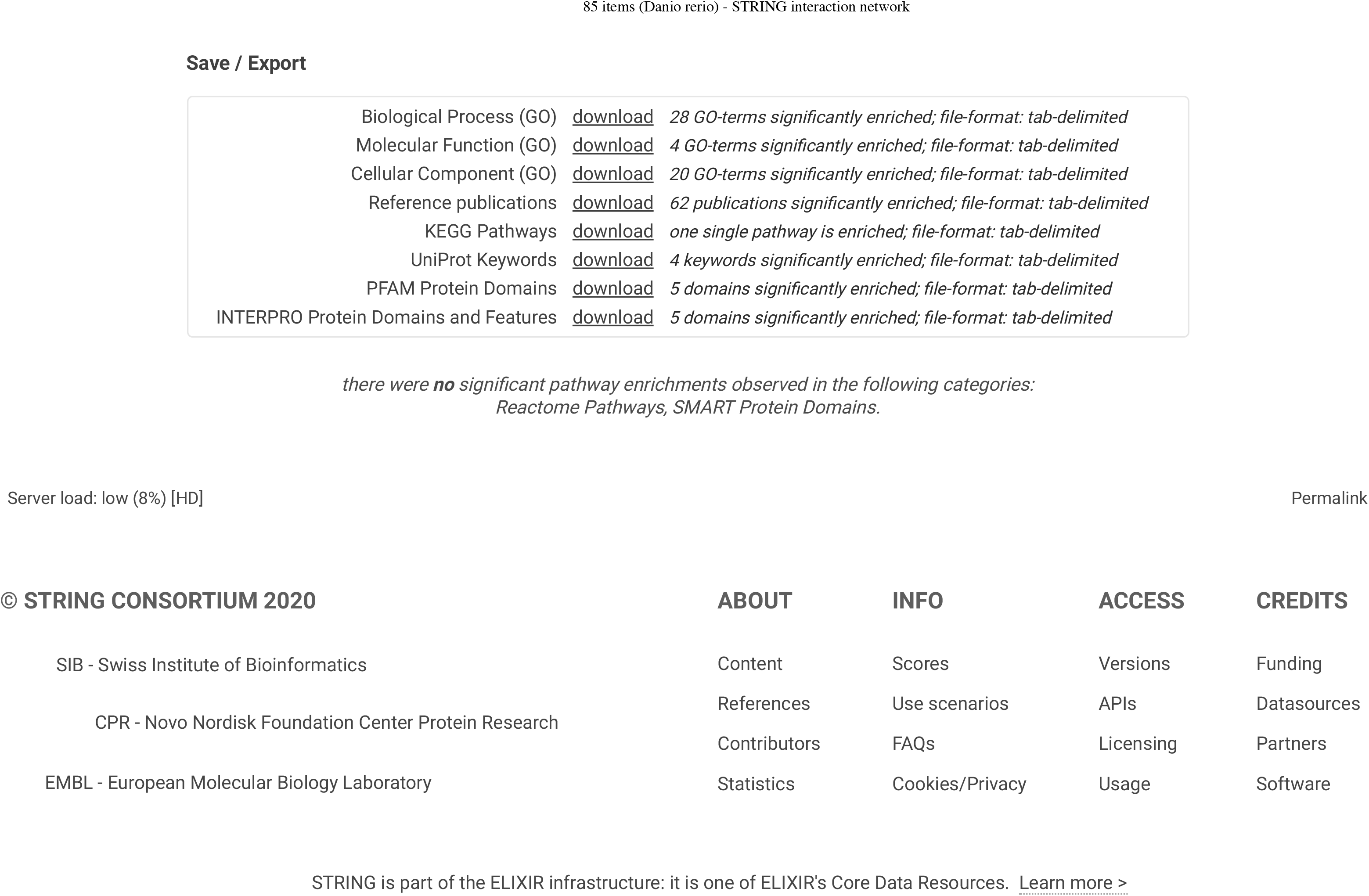

|  |  |
| --- | --- |
| number of nodes: 85 | expected number of edges: 22 |
| number of edges: 38 | PPI enrichment p-value: 0.000941 |
| average node degree: 0.894 | *your network has significantly more interactions* |
| avg. local clustering coefficient: 0.373 | *than expected (<u>what does that mean?</u>)* |

Functional enrichments in your network
| Biological Process (GO) |  |  |  |
| --- | --- | --- | --- |
| *GO-term* | *description* | *count in gene set* | *false discovery rate* |
| GO:0090304 | nucleic acid metabolic process | 10 of 689 | 0.0133 |
| GO:0044237 | cellular metabolic process | 14 of 1406 | 0.0133 |
| GO:0033554 | cellular response to stress | 5 of 145 | 0.0133 |
| GO:0009987 | cellular process | 19 of 2089 | 0.0133 |
| GO:0007006 | mitochondrial membrane organization | 3 of 15 | 0.0133 |
| GO:0007005 | mitochondrion organization | 4 of 68 | 0.0133 |
| GO:0006839 | mitochondrial transport | 3 of 37 | 0.0133 |
| GO:0006807 | nitrogen compound metabolic process | 13 of 1299 | 0.0133 |
| GO:0006139 | nucleobase-containing compound metabolic process | 11 of 771 | 0.0133 |
| GO:0006974 | cellular response to DNA damage stimulus | 4 of 100 | 0.0134 |
| GO:0044260 | cellular macromolecule metabolic process | 11 of 1006 | 0.0138 |
| GO:0044238 | primary metabolic process | 13 of 1373 | 0.0138 |
| GO:0043170 | macromolecule metabolic process | 12 of 1174 | 0.0138 |
| GO:0017004 | cytochrome complex assembly | 2 of 9 | 0.0153 |
| GO:0071704 | organic substance metabolic process | 13 of 1423 | 0.0168 |
| GO:0010842 | retina layer formation | 2 of 10 | 0.0168 |
| GO:0007007 | inner mitochondrial membrane organization | 2 of 10 | 0.0168 |
| GO:0097190 | apoptotic signaling pathway | 2 of 14 | 0.0273 |
| GO:0033108 | mitochondrial respiratory chain complex assembly | 2 of 15 | 0.0284 |
| GO:0065007 | biological regulation | 12 of 1366 | 0.0292 |
| GO:0048592 | eye morphogenesis | 3 of 66 | 0.0292 |
| GO:0051716 | cellular response to stimulus | 7 of 522 | 0.0296 |
| GO:0050896 | response to stimulus | 8 of 683 | 0.0306 |
| GO:0006281 | DNA repair | 3 of 72 | 0.0311 |
| GO:0071840 | cellular component organization or biogenesis | 8 of 726 | 0.0407 |
| GO:0050789 | regulation of biological process | 11 of 1272 | 0.0419 |
| GO:0048646 | anatomical structure formation involved in morphogenesis | 4 of 183 | 0.0485 |
| GO:0009790 | embryo development | 5 of 303 | 0.0485 |
| *(less ...)* |  |  |  |

| Molecular Function (GO) |  |  |  |
| --- | --- | --- | --- |
| *GO-term* | *description* | *count in gene set* | *false discovery rate* |
| GO:0005488 | binding | 18 of 1569 | 0.00029 |
| GO:0003677 | DNA binding | 7 of 372 | 0.0139 |
| GO:0003684 | damaged DNA binding | 2 of 7 | 0.0153 |
| GO:0043167 | ion binding | 9 of 832 | 0.0474 |

| Cellular Component (GO) |  |  |  |
| --- | --- | --- | --- |
| *GO-term* | *description* | *count in gene set* | *false discovery rate* |
| GO:0043231 | intracellular membrane-bounded organelle | 21 of 1613 | 3.43e-06 |
| GO:0005739 | mitochondrion | 7 of 257 | 0.00027 |
| GO:0044429 | mitochondrial part | 6 of 179 | 0.00031 |
| GO:0031966 | mitochondrial membrane | 5 of 134 | 0.00082 |
| GO:0044446 | intracellular organelle part | 12 of 1024 | 0.00098 |
| GO:0005634 | nucleus | 12 of 1043 | 0.0010 |
| GO:0044455 | mitochondrial membrane part | 3 of 34 | 0.0014 |
| GO:0032991 | protein-containing complex | 9 of 631 | 0.0014 |
| GO:0044444 | cytoplasmic part | 10 of 1002 | 0.0077 |
| GO:0032592 | integral component of mitochondrial membrane | 2 of 16 | 0.0077 |
| GO:0070013 | intracellular organelle lumen | 5 of 304 | 0.0140 |
| GO:0005743 | mitochondrial inner membrane | 3 of 87 | 0.0140 |
| GO:0005737 | cytoplasm | 12 of 1534 | 0.0146 |
| GO:0044428 | nuclear part | 5 of 326 | 0.0155 |
| GO:0031090 | organelle membrane | 6 of 489 | 0.0182 |
| GO:0044425 | membrane part | 8 of 844 | 0.0202 |
| GO:0005741 | mitochondrial outer membrane | 2 of 34 | 0.0202 |
| GO:0031461 | cullin-RING ubiquitin ligase complex | 2 of 36 | 0.0211 |
| GO:0005654 | nucleoplasm | 3 of 140 | 0.0321 |
| GO:0098805 | whole membrane | 3 of 154 | 0.0390 |

(less ...)
| Reference publications |  |  |  |
| --- | --- | --- | --- |
| publication | (year) title | count in gene set | false discovery rate |
| PMID:20830285 | (2010) Thyrotroph embryonic factor regulates light-induced... | 4 of 22 | 0.0033 |
| PMID:21390203 | (2011) The light responsive transcriptome of the zebrafish: ... | 5 of 75 | 0.0072 |
| PMID:25079074 | (2014) Redox state and mitochondrial respiratory chain fun... | 3 of 17 | 0.0175 |
| PMID:21756345 | (2011) Cellular expression of Smarca4 (Brg1)-regulated ge... | 4 of 47 | 0.0175 |
| PMID:20415721 | (2010) Cell resilience in species life spans: a link to inflam... | 3 of 15 | 0.0175 |

(more ...)
| KEGG Pathways |  |  |  |
| --- | --- | --- | --- |
| pathway | description | count in gene set | false discovery rate |
| dre04340 | Hedgehog signaling pathway | 3 of 52 | 0.0281 |

| UniProt Keywords |  |  |  |
| --- | --- | --- | --- |
| keyword | description | count in gene set | false discovery rate |
| KW-0539 | Nucleus | 16 of 1851 | 0.0262 |
| KW-0802 | TPR repeat | 4 of 114 | 0.0289 |
| KW-0496 | Mitochondrion | 6 of 319 | 0.0289 |
| KW-0677 | Repeat | 14 of 1754 | 0.0358 |

| PFAM Protein Domains |  |  |  |
| --- | --- | --- | --- |
| domain | description | count in gene set | false discovery rate |
| PF13844 | Glycosyl transferase family 41 | 2 of 2 | 0.0098 |
| PF07719 | Tetratricopeptide repeat | 3 of 31 | 0.0098 |
| PF00515 | Tetratricopeptide repeat | 3 of 27 | 0.0098 |
| PF13431 | Tetratricopeptide repeat | 2 of 6 | 0.0113 |
| PF13414 | TPR repeat | 2 of 13 | 0.0334 |

| INTERPRO Protein Domains and Features |  |  |  |
| --- | --- | --- | --- |
| domain | description | count in gene set | false discovery rate |
| IPR001440 | Tetratricopeptide repeat 1 | 3 of 14 | 0.0056 |
| IPR037919 | UDP-N-acetylglucosamine-peptide N-acetylglucosaminyltr... | 2 of 2 | 0.0082 |
| IPR029489 | O-GlcNAc transferase, C-terminal | 2 of 2 | 0.0082 |
| IPR019734 | Tetratricopeptide repeat | 4 of 121 | 0.0478 |
| IPR013026 | Tetratricopeptide repeat-containing domain | 4 of 121 | 0.0478 |

|  |  |  |
| --- | --- | --- |
| Biological Process (GO) | download | 28 GO-terms significantly enriched; file-format: tab-delimited |
| Molecular Function (GO) | download | 4 GO-terms significantly enriched; file-format: tab-delimited |
| Cellular Component (GO) | download | 20 GO-terms significantly enriched; file-format: tab-delimited |
| Reference publications | download | 62 publications significantly enriched; file-format: tab-delimited |
| KEGG Pathways | download | one single pathway is enriched; file-format: tab-delimited |
| UniProt Keywords | download | 4 keywords significantly enriched; file-format: tab-delimited |
| PFAM Protein Domains | download | 5 domains significantly enriched; file-format: tab-delimited |
| INTERPRO Protein Domains and Features | download | 5 domains significantly enriched; file-format: tab-delimited |

there were **no** significant pathway enrichments observed in the following categories: Reactome Pathways, SMART Protein Domains.
Server load: low (8%) [HD]
Permalink
© STRING CONSORTIUM 2020
ABOUT
INFO
ACCESS
CREDITS
SIB - Swiss Institute of Bioinformatics

## Notes

### Competing Interest Statement

The authors have declared no competing interest.

